# A collection of designed peptides to target SARS-Cov-2 – ACE2 interaction: Pep*I*-Covid19 database

**DOI:** 10.1101/2020.04.28.051789

**Authors:** Ruben Molina, Baldo Oliva, Narcis Fernandez-Fuentes

**Affiliations:** Structural Bioinformatics Lab, Department of Experimental and Health Science, Universitat Pompeu Fabra, Barcelona 08003, Catalonia, Spain; Department of Biosciences, U Science Tech, Universitat de Vic-Universitat Central de Catalunya, Vic 08500, Catalonia, Spain; Institute of Biological, Environmental and Rural Sciences, Aberystwyth University, Aberystwyth SY23 3EB, United Kingdom

## Abstract

The angiotensin-converting enzyme 2 is the cellular receptor used by SARS coronavirus SARS-CoV and SARS-CoV-2 to enter the cell. Both coronavirus use the receptor-binding domain (RBD) of their viral spike protein to interact with ACE2. The structural basis of these interactions are already known, forming a dimer of ACE2 with a trimer of the spike protein, opening the door to target them to prevent the infection. Here we present Pep*I-*Cov19 database, a repository of peptides designed to target the interaction between the RDB of SARS-CoV-2 and ACE2 as well as the dimerization of ACE2 monomers. The peptides were modelled using our method PiPreD that uses native elements of the interaction between the targeted protein and cognate partner that are subsequently included in the designed peptides. These peptides recapitulate stretches of residues present in the native interface plus novel and highly diverse conformations that preserve the key interactions on the interface. Pep*I*-Covid19 database provides an easy and convenient access to this wealth of information to the scientific community with the view of maximizing its potential impact in the development of novel therapeutic agents.

## Introduction

In December 2019, a disease affecting predominantly the respiratory system emerged in Wuhan, province Hubei, China, caused by a new coronavirus: SARS-CoV-2. The SARS-CoV-2 virus is closely related to SARS-CoV virus responsible for an outbreak in 2002^1^. To infect the cells, both SARS-CoV-2 and SARS-CoV use the angiotensin-converting enzyme 2 (ACE2) as a keyhole, binding it via the receptor-binding domain (RDB) of the spike protein (S protein) together serine protease TMPRSS^2–5^. This interaction is a key step in viral infection of the cells and thus, preventing this association the viral charge arises as a valid therapeutic strategy.

The inhibition of protein-protein interactions (PPIs) using peptides is a valid approach that have been gaining traction in recent years given the limitation of traditional small, drug-like, chemicals to target interfaces^6^. Particularly in the context of viral infection examples such as the FDA-approved peptide Enfurvirtide^7^ and further research (reviewed in ^8^) justify the use of peptides as potential therapeutic agents to block PPIs.

The structural details of the interaction between ACE2 and the RBD of the S protein, both for the SARS-CoV^2^ and SARS-CoV-2^4, 9^ viruses are known. The structural information of these protein complexes can be used by programs such the one developed by us, PiPreD^10^, to guide the modelling and design orthosteric peptides to target protein interfaces of interest. We have therefore leveraged on the existing structural information available on the interaction between RBD and ACE2 to model and design peptides targeting two relevant interactions for the complex: 1) the interaction between RBD and ACE2; and 2) the interaction between ACE2 monomers.

We have compiled this information in a public repository: Pep*I*-Covid19 to make it available to the scientific community, particularly those scientists and groups searching for novel therapeutic agents. Besides the sequences included in repository, the modelled structures of the protein-peptides complexes can represent also the starting point for further refinement and redesign using alternative approaches that can yields novel peptide sequences. Finally, we aim at keeping Pep*I*-Cov19 an up-to-date and alive resource, and thus any new structural data on protein complexes related to covid19 will be duly processed and included in the repository.

## Material and Methods

### Surface targeted and design of peptides

We use the structure of the SARS-CoV-2 spike receptor-binding domain bound with ACE2 (PDB code 6m0j) ^9^ and SARS-CoV-2 spike receptor-binding domain bound to full length ACE2 (PDB code 6m17)^4^. The two protein complexes, 6m0j and 6m17, were used in the case of the interface mediating the interaction between the spike protein (S-prot) and ACE2 while the latter was use to model and design peptides to target the interface between ACE2 monomers (Figure 1).

**Figure 1.**
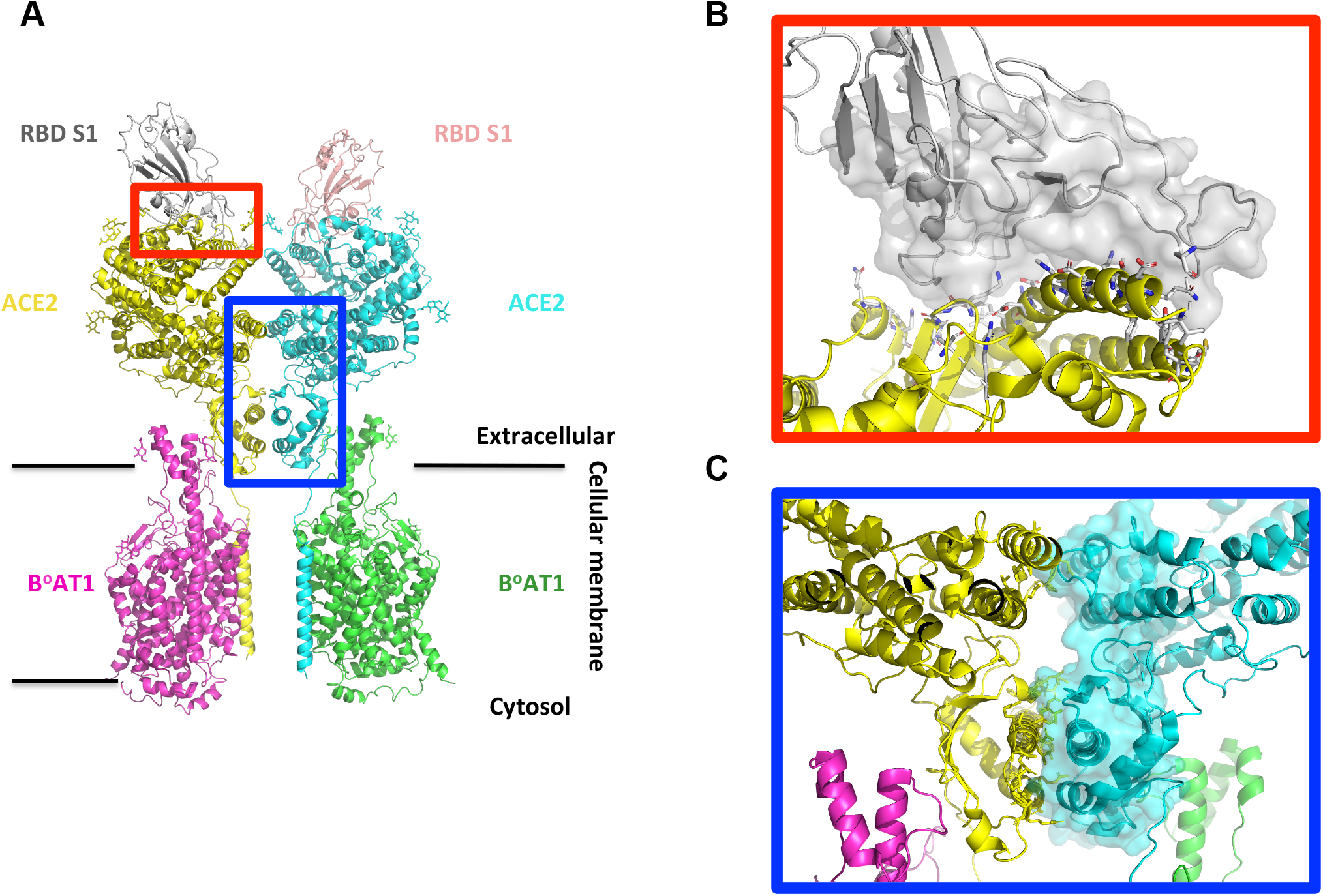
Interface targeted by designed peptides. (A) Cartoon representation of full length human ACE2 (yellow, cyan) / BoAT1 (magenta, green) and RDB S-prot (grey, light brown); extracellular and membrane embedded regions are shown. (B) Interface A. Detail of the interface between ACE2 (yellow, cartoon) and RDB Sprot (grey, surface). (C) Interface D. Detail of the interface between ACE2 monomers.

Peptides were modelled and designed using PiPreD and Rosetta^11^ as previously described^10^. The modelling step relies on a library of iMotifs that are fitted in the interface to target using the so called anchor residues. The parameters used in the modelling stage were: (i) a maximum distance between Cα iMotifs-anchors of 0.5 Ang., and (ii) a root mean square deviation value smaller than 1.0 Ang. upon structural superposition iMotifs-anchors. Once the iMotifs were structurally fitted and prior to the designing stage, peptides with low interface packing and mimicking less than 4 anchor residues were discarded.

The design of peptides was done using the backrub application ^12^ within the Rosetta suite^11^. The backrub motions allow the limited movement of the main-chain of the peptide. It optimizes the interactions with the interface while allowing the design of any position not structurally aligned with anchor residues, where residue type is preserved although rotameric changes were allowed. The residues on the interface belonging to the protein were also allowed to repack. Once the designing stage was completed, the FlexPepDock application^13^ in “score mode only” was used to obtain severals scores (i.e. interface score or peptide score) and other measures such as buried surface are of the interface between peptide and protein

### Database design, implementation and interfacing

Information about the designed peptides is stored in an SQLite3 database. The web interface accessible for the user is generated with the Flask Python Microframework. The structure of the complex protein-peptides is rendered with PV, a WebGL-based JavaScript API (https://biasmv.github.io/pv/).

## Results and Discussion

### Interface S1 RDB-ACE2 and ACE2/ACE2

Two different interfaces were considered in this study: the interface between the RBD of viral S protein, henceforth referred to as interface A, and ACE2 and the extracellular region of the interface mediating the dimerization of ACE2 protein, interface D (Figure 1).

For interface A, peptides were designed to target the surface of the viral protein, i.e. the RBD of the S-prot, hence the anchor residues were derived from the residues of ACE2 protein. The area of the interface is around 800 Ang^2^ comprising 26 anchor residues. The interaction between both proteins is dominated by extensive contacts between loops of RDB and the a-1 (H1) helix of ACE2 (Figure 1 panel B). The modelling step prior to designing stage yielded almost 15M peptide, which after discarding poorly packed peptides and those mimicking less than 3 anchor residues resulted in a total of number of 450,000 peptides were taken forward to the designing stage.

For interface D, the number of anchor residues was 37, spanning an interface with a surface substantially larger than interface A (approximately 2200 Ang^2^.) The structural elements mediating the interactions between both monomers are mainly stacking mirroring helices from both monomers. As discussed in the publication describing the structure^4^, ACE2 dimerizes through two different interfaces involving the peptidase (PD) and Neck domains. In the central region, between both interfaces, there are not direct interactions between ACE2 monomers; however, it was also considered for the modelling of peptides. Interestingly, peptides spanning the two distant interfaces include native elements of the interface (i.e. mimicry of anchors residues) but also *de novo* interactions (Figure 1, panel C, surface representation). The modelling step yielded over 26.5 M peptides. Upon discarding peptides with a poor interface packing and structurally aligning with less than 3 anchor residues the number dropped to over 1.5M.

### Native elements of the interfaces are recapitulated by designed peptides

As discussed, PiPreD relies on native elements of the interface to target in the form of disembodied interface residues, anchor residues, although without considering the connectivity between them. Thus, the resulting peptides are not mere regions or elements of the interface spliced from the native complex. Nonetheless, designed peptides also recapitulate these native elements of the interface. Examples are shown in Figure 2 (panels A and B), for each explored interface. In the case of interface A, different peptides recapitulated the main H1 helix (partial or entire) with minor conformation adjustments. Likewise, in interface D two peptides show a helical conformation similar to the helix that mediates the interaction between ACE2 monomers.

**Figure 2.**
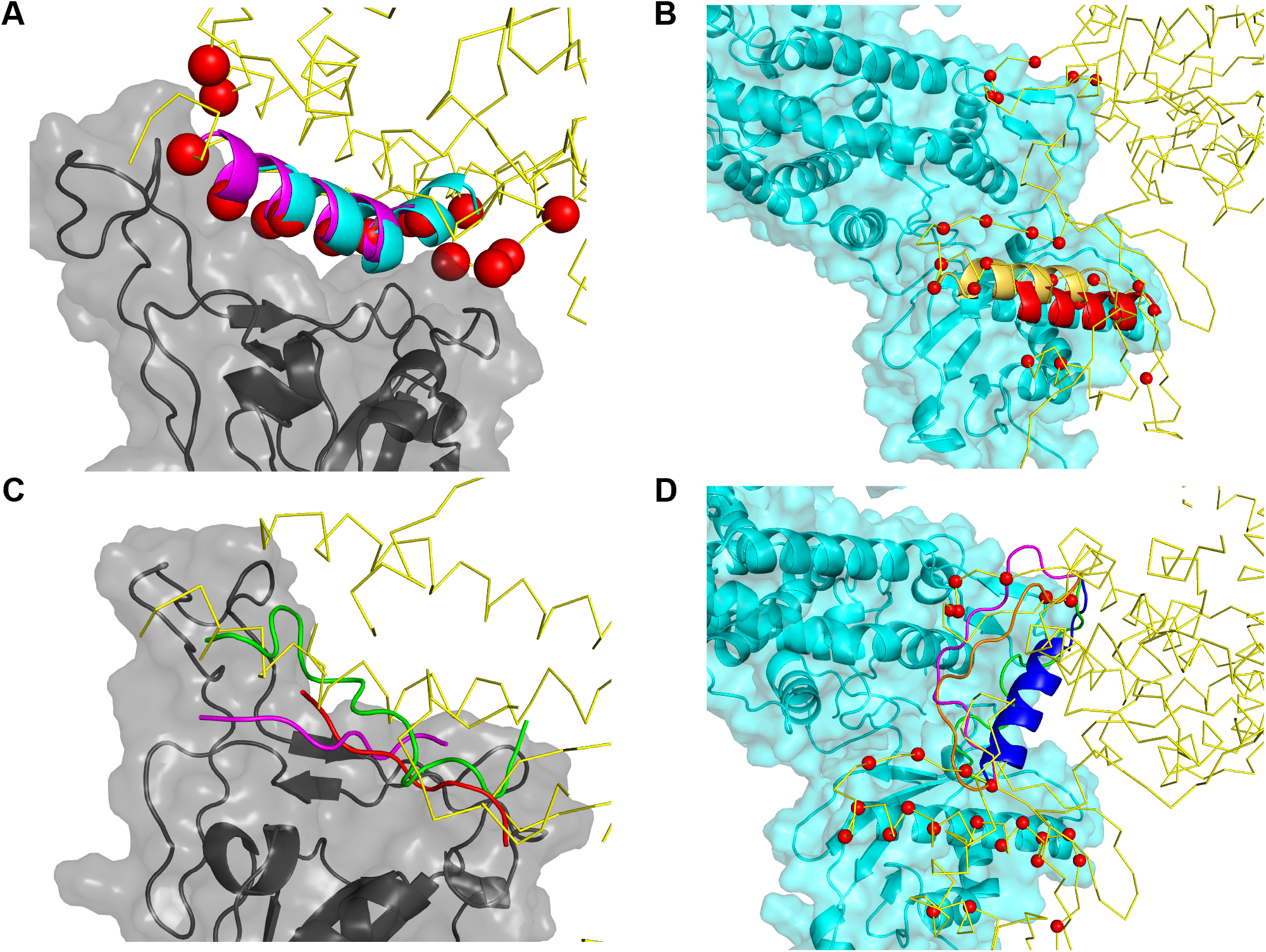
Structural model of the complexes RBD S-prot and ACE2 bound to peptides. Panels A and C shows the structural model of several peptides of the designed peptides bound to interface A recapitulating native elements of the interface (A) or new conformation (C). The RBD of Sprot is shown in grey and cartoon and surface representation while ACE2 is shown as traces of Ca; anchor residues a shown as red spheres. Likewise panels B and D show the structural models of peptides bound to interface D also recapitulating native elements (B) or new conformations (D).

### Structurally diversity among designed peptides

PiPreD provided peptides with conformations unobserved in the native complex. These novel conformations incorporated native elements of the interface in the form of anchor residues, mimicking or surrogating them, but also introducing novel interactions between the peptides and the targeted surface.

For interface A, the RDB of S-prot binds to ACE2 primarily through a long alpha helix (H1) and to a lesser extend by a second, shorter, helix. However, over half of the total number of peptides designed to target the surface of RDB have an extended, linear, conformation. Three examples are shown Figure 2 (panel C) with peptides presenting an extended conformation mimicking the interactions of the main H1 helix.

Peptides targeting interface D also present conformations unobserved in the native interface. Interestingly, the association between ACE2 is mediated by two different interfaces; one involving the PD domain and the other close to the membrane, the Neck domain. Peptides bridging between the two surfaces apart were designed both in helical and extended conformations (Figure 2, panel C), potentially increasing the affinity of the peptides through a synergistic effect rather than the sum of each single interface.

### PepI-Covid19 database repository: access and functionalities

An online repository to the peptides designed is freely accessible at the Pep*I*- Covid19 database (http://aleph.upf.edu/pepicovid19/). The repository is web interfaced allowing user to perform queries based on parameters such as the conformation of the peptides, size and predicted binding energy as per the Rosetta energy score^11^. Users can also query the information based on a number of interface statistics such the surface area of the interface between protein and peptides (in Ang^2^), the number of hydrogen bonds and unsatisfied hydrogen bonds (in case of buried donor or acceptors groups) at the interface, packing^14^ or the peptide score.

The query returns a list of the peptides that fulfill the conditions set on the query. The resulting list can then filter by primary amino acid sequence, and/or sorted by query parameters (e.g. size). Upon selecting a specific peptide, a page with specific information on the peptide is shown together with an applet allowing the three dimensional visualization of the protein-peptide structural complex (Figure 3.)

**Figure 3.**
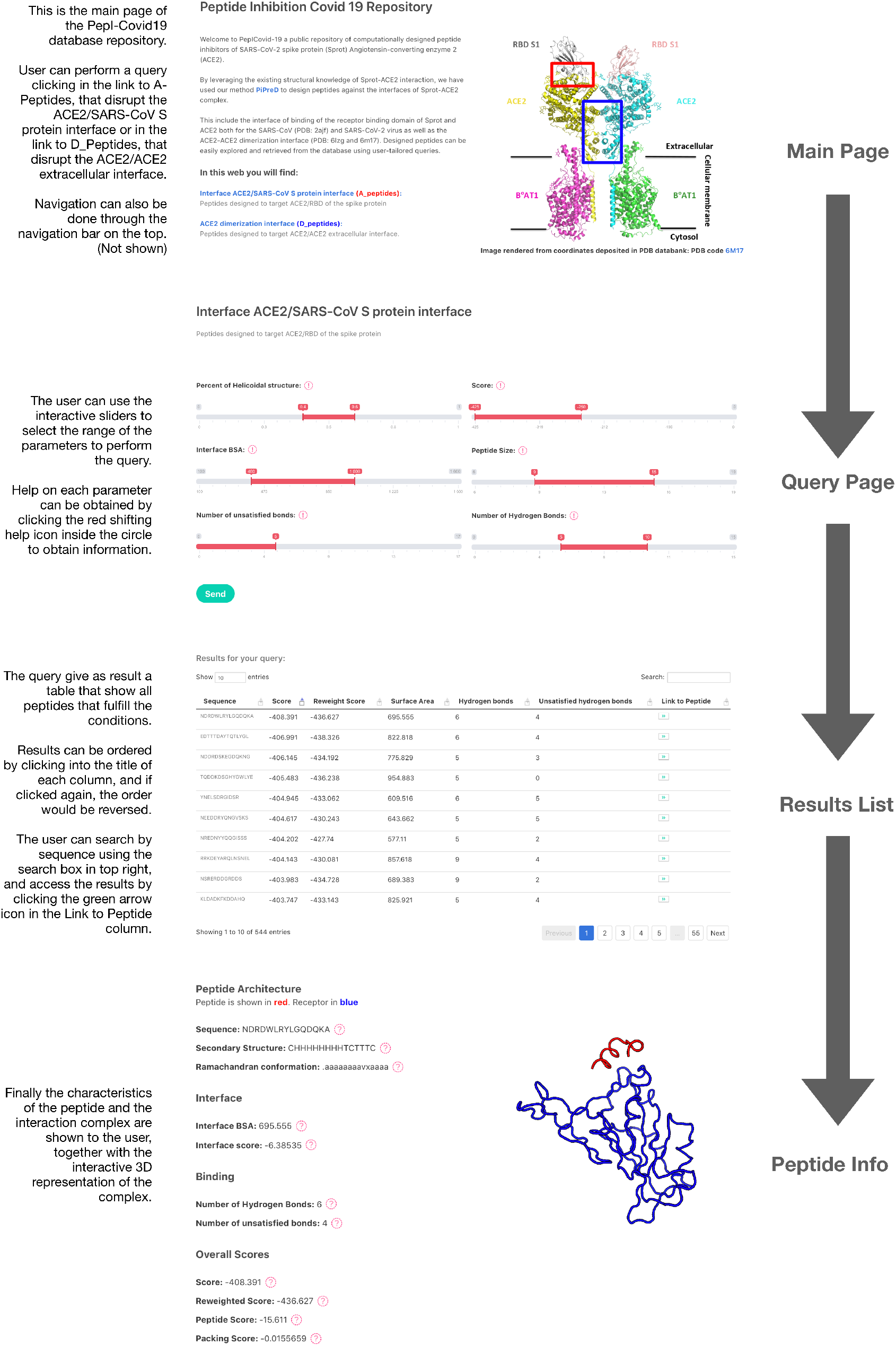
Pep*I-*Covid19 database web interface. From top to bottom the different web-pages shown in the repository that allow users to query, search and filter results as well as visualize the three-dimensional structure of protein complex interactively.

## Conclusion

Here we present a repository of designed peptides to interfere in the early stages of invasion process by SARS-CoV-2. These peptides present a large structural sampling, targeting both the RBD/ACE2 as well as the extracellular ACE2/ACE2 interface. The repository allows for tailored queries and searches and a convenient retrieval of information. Users can visualize the structural model of the protein-peptide interactively. We believe the information included in Pep*I-*Cov19 database is a significant and worth asset to current efforts devoted to find novel therapeutic agents and strategies to fight SARS-CoV-2. Indeed, the information of protein sequences of potential peptide inhibitors for experimental guidance or structural data of peptides as basis for further computational work represent two of the elements where Pep*I*-Cov19 database can contribute to such efforts. Moreover, the repository will be updated as structural information becomes available on protein complexes related to SARS-CoV-2 cell infection.

## Acknowledgements

Authors thank the research groups and members within that solved the structures used in this work making it possible. Authors thank the technical support provided by the members of the GRIB IT team: Alfons Gonzalez-Pauner and Miguel A. Sanchez Gomez.

